# AlphaFold-assisted design of a modular Laminin E8 Fragment Panel

**DOI:** 10.64898/2026.11.20.768189

**Authors:** Lauren Litzau, Andrew C. Kruse, Stephen C. Blacklow

## Abstract

Laminins are large, heterotrimeric glycoproteins that are a major structural component of basement membranes and the extracellular matrix. Although they are difficult to produce recombinantly because of their large size, a smaller fragment of the laminin heterotrimer that supports cell adhesion and cell migration, termed E8, can be recombinantly expressed. Here, we use AlphaFold to optimize laminin E8 fragments and design a panel of interchangeable laminin α, β, and γ chains that can be combined to produce all isoforms recombinantly. These recombinant E8 fragments support adhesion of cells bearing their cognate integrins, and will be useful tools for stem-cell, organoid, and laminin-binding integrin biology research.

**Impact statement:** Laminins are large, heterotrimeric proteins that are difficult to express recombinantly, because of their large size. This work reports a laminin E8 fragment design optimized by AlphaFold and a modular expression pipeline for the study of all laminin isoforms. Laminin E8 fragments are useful reagents for studying laminin-binding integrin interactions and for stem-cell and organoid biology protocols.

## Introduction

Laminins are the major extracellular matrix protein in the basement membrane. They are bound by cell adhesion molecules and receptors including integrins and dystroglycan (1). They are the first extracellular matrix (ECM) proteins expressed in embryos and are important for embryonic development and tissue structural integrity (2). Mutations in laminins are commonly embryonic lethal, underlining their critical role in development (3). In the stem-cell biology field, laminins are used in induced pluripotent stem cell (iPSC) and organoid differentiation protocols (4).

Laminins are large, 400 to 900 kDa, glycoproteins composed of an α, β, and γ chain. Laminin nomenclature defines each heterotrimer isoform based on the identity of its α, β, and γ subunints. For example, laminin-332 (Lm-332) contains the α3, β3, and γ2 isoforms as its three subunits (5). Laminins have 4 “arms” that are capped by globular domains **(Figure 1A)** (6). The long arm is formed by a heterotrimeric coiled-coil, which is stabilized by four disulfides - three at the N-terminus and one at the C-terminus between the β and γ subunits. The end of the coiled- coil is capped by laminin globular-like (LG) repeats of the α chain, with the three other short arms each derived from a single chain, α, β, or γ (7).

**Figure 1.**
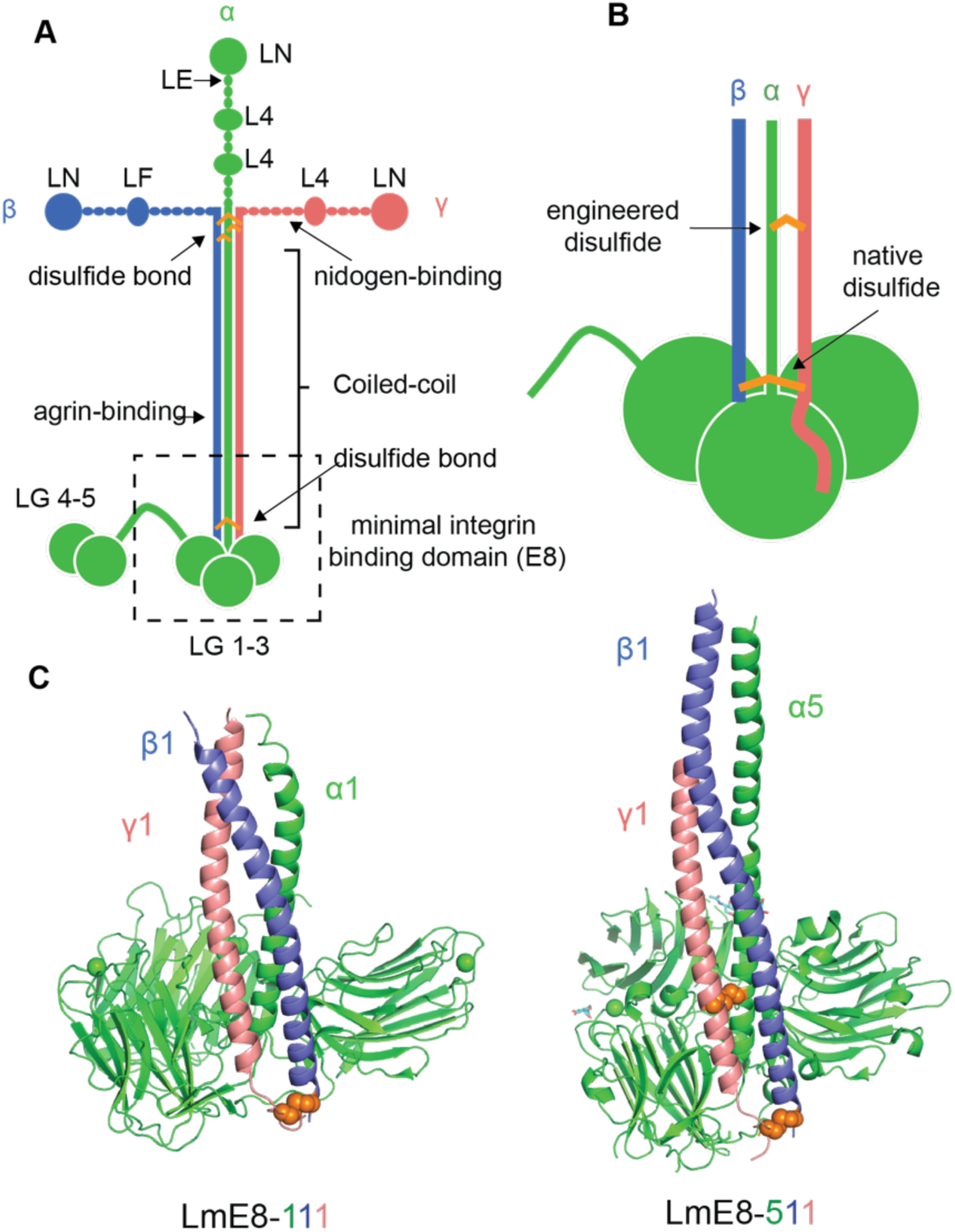
Laminin heterotrimer and the E8 fragment laminin binding. **(A)** Schematic of a representative laminin heterotrimer, showing the various modular elements present on each chain. **(B)** Schematic of the truncated LmE8 structure indicating the location of the native engineered disulfides. Adapted from (7). **(C)** Cartoon rendering of the x-ray crystal structures of the LmE8-511 (PDB: 5XAU) and LmE8-111 (PBD: 5MC9) fragments (14,15). Interchain disulfide bonds are shown using orange spheres.

Biochemical studies have established that laminins form multivalent assemblies with other laminins through their laminin N-terminal (LN) domains (8,9), bind agrin through the coiled-coil (10), and bind nidogen through the fourth laminin-type EGF-like repeat (LE4) of the γ1 subunit (11). Importantly, the C-terminal coiled-coil arm interacts with integrin and dystroglycan adhesion receptors (12).

Laminins are large and difficult to produce recombinantly. To purify full length laminin, isoforms are commonly immunoaffinity purified from endogenous sources: mouse Engelbreth- hold-Swarm tumor tissues, human placenta, human gastric cancer cells, human glioma cells, and human lung adenocarcinoma cells for Lm-111, Lm-211/221, Lm-332, Lm-411, and Lm-511/521 respectively (13). However, a fragment of laminin supports cell migration and integrin binding (12). This fragment, termed the Laminin E8 fragment (LmE8), contains the C-terminal region of the long arm coiled-coil, and LG repeats 1-3 **(Figure 1B)**. The E8 fragments are smaller, ∼100kDa, and can be recombinantly expressed and purified. For structural biology and x-ray crystallography efforts, an engineered disulfide was added between the α and γ coiled-coil to further stabilize the heterotrimer and mimic the N-terminal coiled-coil disulfides (14). Structures of LmE8-111 and LmE8-511 were solved by x-ray crystallography and cryo-electron microscopy **(Figure 1C)** (14,15). A structure of LmE8-511 bound to the laminin-binding integrin adhesion receptor α6β1 revealed that the LG domains interact with the integrin headpiece or ligand binding domain, and a glutamate residue in the γ chain C-terminal tail coordinates a metal ion to induce integrin conformational change and activate signaling (16).

Here, we used AlphaFold3 to optimize laminin E8 construct designs for all laminin isoforms (17). In humans, there are 17 biochemically confirmed laminin isoforms composed of 5 α, 4 β, and 3 γ chains. We took inspiration from solved laminin structures, used AlphaFold3 to align all coiled-coils and remove unstructured loops, and tested whether adding an engineered disulfide between the α and γ coiled-coil increases the efficiency of protein expression. We found that coiled-coiled truncation, unstructured loop truncation, and engineered disulfide removal increased laminin E8 fragment yield. The panel of laminin subunits that we have designed are modular and can be combined with one another to express all human laminin E8 isoforms.

## Results

### Initial LmE8 fragment design with an engineered disulfide bond

Our initial attempt to design a complete LmE8 panel was based on the strategy employed to make LmE8-511 for structure determination (14). This strategy introduces an engineered disulfide between the α5 and γ1 chains in the coiled coil. We applied the disulfide strategy to other chains and determined chain length using sequence alignments with LmE8-511 as a template. Our panel includes laminin subunits: α1, α2, α3, α4, α5, β1, β2, β3, γ1, γ2, and γ3. We did not include β4 in the panel, because β4 was previously thought to be a pseudogene and its expression was only recently confirmed by immunohistochemistry in the colon (18,19).

The sequences of the coiled-coil regions of all chains aligned well, and engineered disulfides were introduced at conserved positions to couple the α and γ chains. Each α chain was modified with an Asp to Cys mutation, and each γ chain received an Ile to Cys mutation **(Figure 2A)**. This engineered disulfide ensures that all three chains are covalently connected because of the native disulfide already present between the β and γ chains. To direct secretion of the expressed protein, constructs were designed with an N-terminal hemagglutinin signal sequence, which was followed by an affinity tag and a GGS linker before the first residue of the mature laminin chain subunit. His6, HA, and Flag tags were used as the affinity handles on α, β, and γ subunits, respectively. Constructs were cloned into the pTarget vector for mammalian expression.

**Figure 2.**
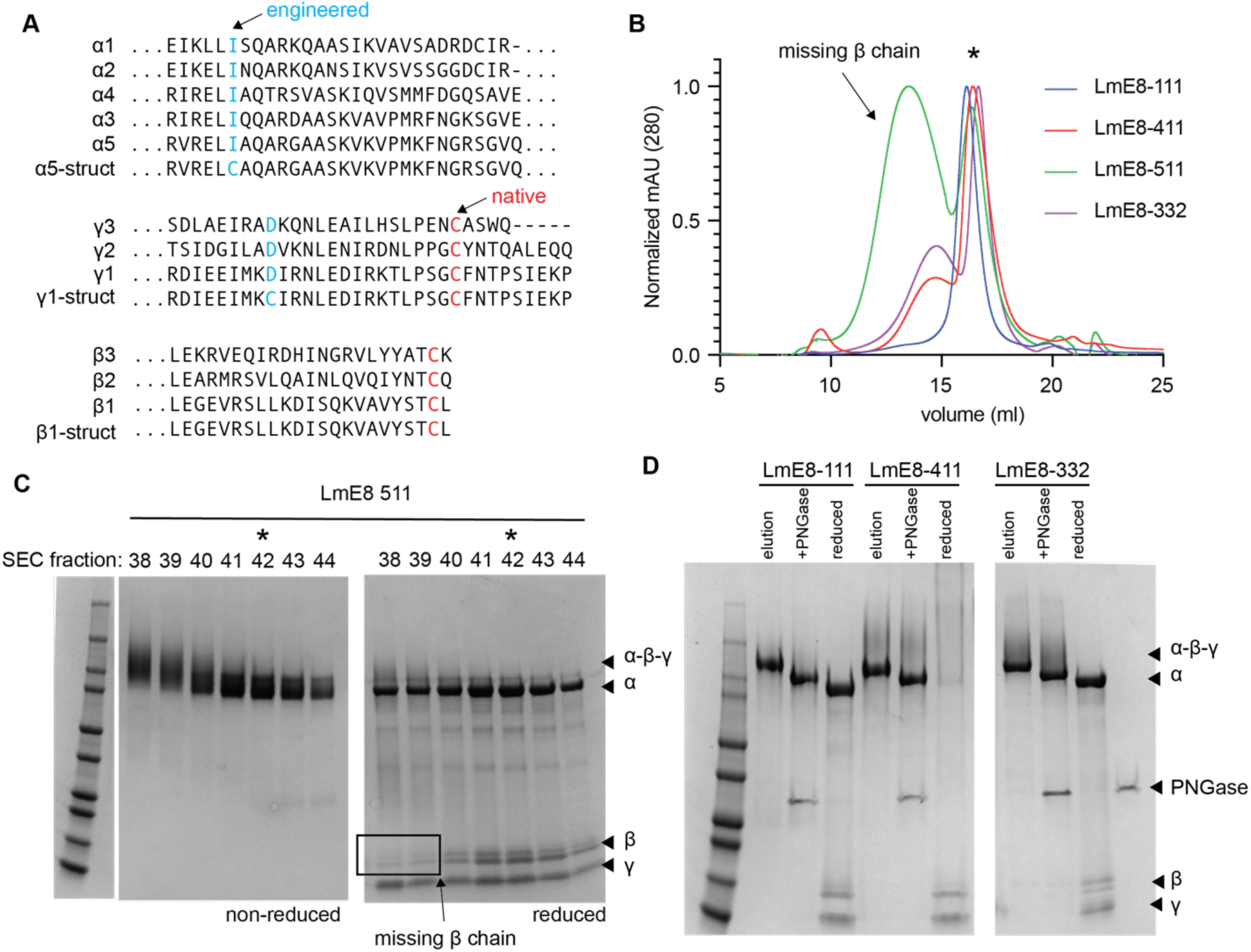
Laminin E8 fragment design with an engineered disulfide bond and protein purification. **(A)** Alignment of α, β, and γ chain coiled-coil domains. Residues mutated to cysteine to form engineered disulfides are shown in blue, and the native cysteines are shown in red. **(B)** Size exclusion chromatography (SEC) traces showing purification of the indicated LmE8 proteins on a Sepharose 6i column. Traces were normalized to set the maximum and minimum 280 nm absorbance values to 1 and 0, respectively. The peak at ∼17 ml (indicated with an asterisk) corresponds to the α-β-γ fragment with all disulfides formed and intact. **(C)** Coomassie-stained gel of LmE8-511 SEC fractions analyzed by SDS-PAGE. Left panel: non-reducing gel; right panel: reducing gel. Fractions 38-40 correspond to the higher molecular weight fraction missing the β chain. The peak with the asterisk (fraction 42) contains all chains. **(D)** Coomassie-stained gels of purified LmE8-111, LmE8-411, and LmE8-332. Lanes compare untreated samples with PNGase-treated and reduced samples, showing that all chains are present, glycosylated, and disulfide linked.

In small scale cultures to test for expression, LmE8-111, LmE8-332, LmE8-411, LmE8- 421, LmE8-521, and LmE8-511 were readily detected and formed the correct disulfides. Purifications were then scaled up for LmE8-332, LmE8-111, LmE8-411, and LmE8-511. After affinity purification, a large fraction of the eluted material contained only alpha and gamma chains **(Figure 2B-C)**, but this impurity could be removed after size-exclusion chromatography. Fractions that were the correct size by size-exclusion chromatography had all chains present, were appropriately glycosylated **(Figure 2D)**, and had formed disulfides that connect all three chains, with one α-β-γ band observed in non-reducing conditions, and individual α, β, and γ bands observed in reducing conditions **(Figure 2D)**.

### LmE8 fragments activate integrin signaling and support cell adhesion

We then evaluated whether the LmE8 fragments were functional, as assessed by integrin binding and signal activation. To probe cell adhesion and integrin signaling, we used a cell-based assay in which we immobilized LmE8 on untreated tissue culture plates or on high binding plates. We then deposited SVG-A fetal astrocyte cells, which express laminin binding integrins ITGα3β1 and ITGα6β1, but not ITGα7β1, onto the plates (ITGα3β1 and ITGα6β1 ectodomains bind to LmE8-332 and Lm-511 with low nanomolar affinity, but do not bind specifically to Lm-111 and Lm-411 (13)). In agreement with the known binding preferences of the ITGα3β1 and ITGα6β1 integrins for their cognate laminins, SVG-A cells adhered most strongly to plates coated with LmE8-332 and LmE8-511 after washing **(Figure 3A)**. Cellular adhesion to these LmE8 fragments also activated FAK phosphorylation, a proximal readout of integrin signaling **(Figure 3B)** (20). Furthermore, in ITGα3β1 knockout SVG-A cells, adhesion to LmE8-332 or LmE8-511 is decreased **(Figure 3C)**, and this adhesion defect is enhanced in ITGα3β1/ITGα6β1 double knockout cells. This outcome is similar to that seen for cell adhesion of ITGα3/ITGα6 KO kidney cells to full length Lm-511 and Lm-332 (21).

**Figure 3.**
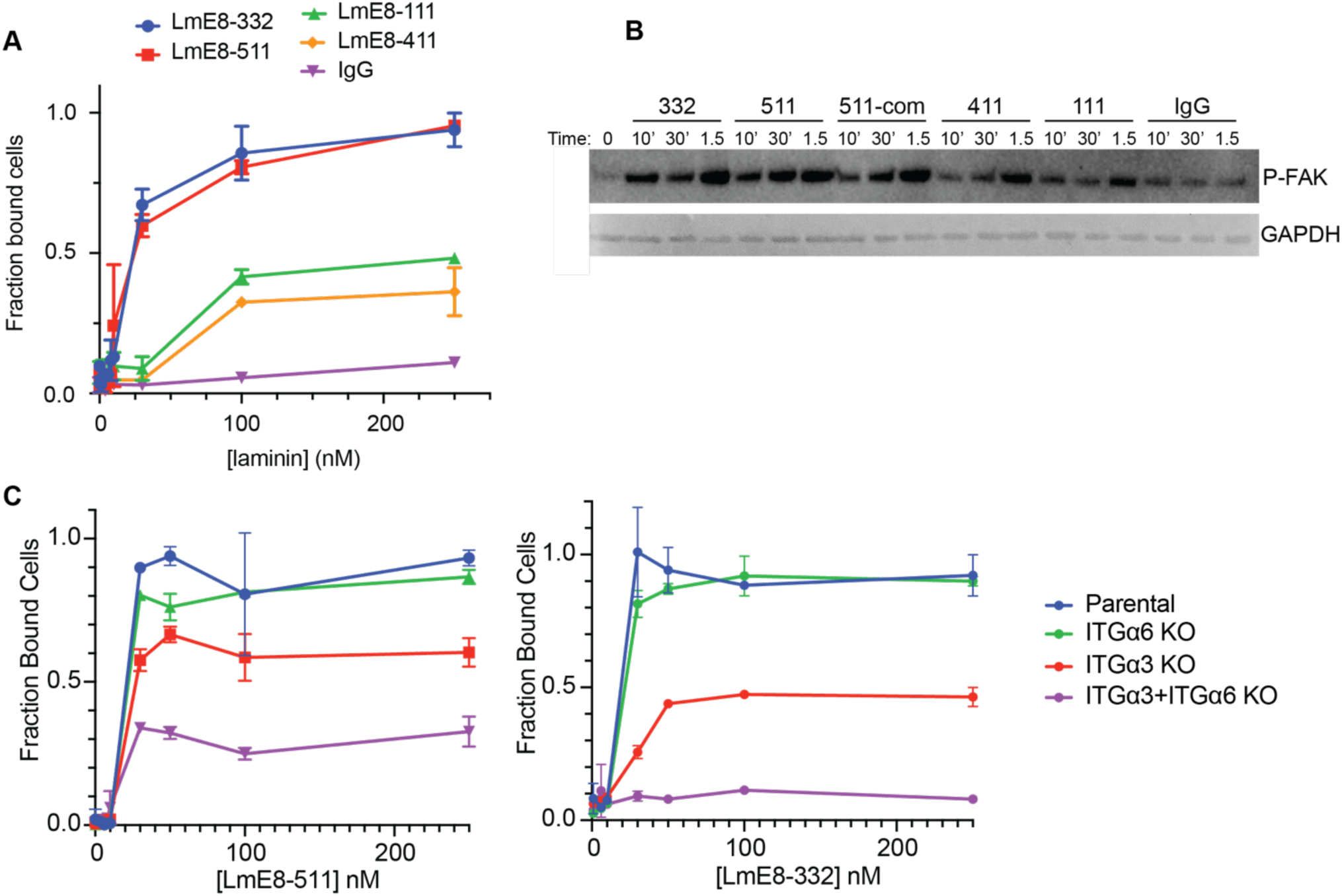
LmE8 fragments are functional in adhesion assays. **(A)** SVG-A cell adhesion assay. The fraction of adherent SVG-A cells is plotted as a function of the concentration of the LmE8 fragment coated on the plate. **(B)** Western blot probing the induction of p-FAK as a function of time after deposition of SVG-A cells onto plates coated with the indicated LmE8 fragments. In- house purified LmE8-511 activates integrin signaling as effectively as the commercial preparation of LmE8-511 (LmE8-511-com). **(C)** SVG-A adhesion assay using integrin knockout cells as indicated. Adhesion independent of laminin-binding integrins on LmE8-511 could be due to the presence of other integrins or adhesion molecules.

These data establish that the designed LmE8 fragments are functional. However, the efficiency of disulfide bond formation and yield of correctly folded product was variable, and these limitations led us to reassess our approach to construct design. For these LmE8 redesign efforts, we utilized AlphaFold3 to improve our LmE8 fragment design length and engineered disulfide positioning, as described below (17).

### AlphaFold assisted LmE8 redesign

We ran AlphaFold3 Multimer predictions for all LmE8 isoforms. In these analyses, the engineered disulfide bonds were predicted to form in the correct arrangement, but an additional unstructured loop after the LG1-3 domains and a coiled-coil helix overhang was also predicted **(Figure 4A)**. As an alternative strategy, we reasoned that we could improve upon the original design by eliminating the engineered disulfides, instead modifying the length of the α chains by truncation to include only structured regions and to optimize the alignment of the coiled-coil segment **(Figure 4B-C).**

**Figure 4.**
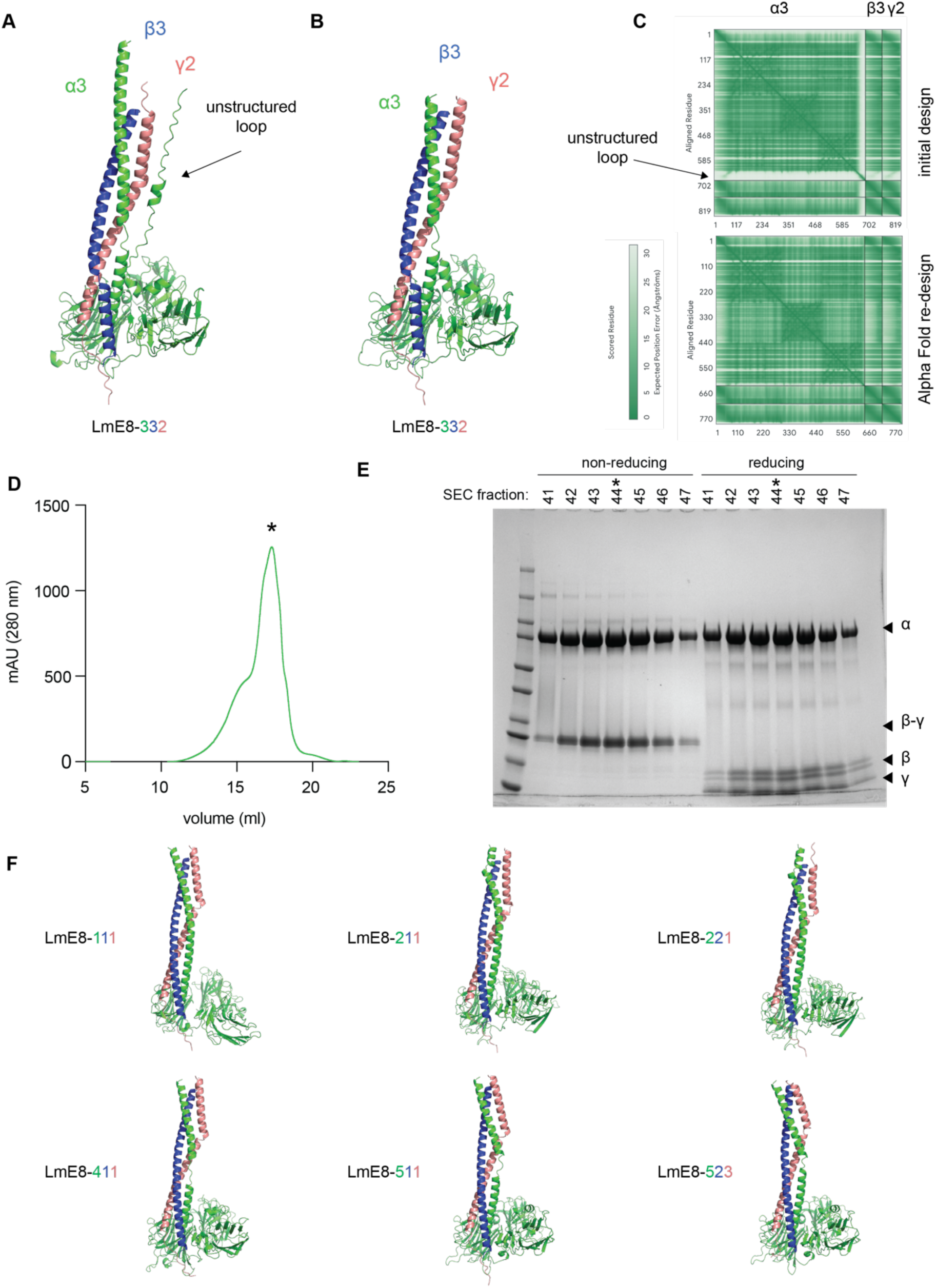
AlphaFold3 assisted design of LmE8 fragments. **(A)** AlphaFold3 prediction of the LmE8-332 structure identified a potential unstructured loop and extended α chain helix, which were removed in the LmE8-332 re-design shown in **(B)**. **(C)** Predicted aligned error plots, which exhibit high confidence predictions for the initial LmE8-332 model (pTM: 0.83, ipTM: 0.82) and the redesigned LmE8-332 construct (pTM: 0.83, ipTM: 0.82). **(D)** Size exclusion chromatography (SEC) trace and of the redesigned LmE8-332 construct on a Superose 6i column. **(E)** Coomassie- stained SDS-PAGE gel of the SEC fractions. The main peak, indicated with an asterisk, contains all chains, with the native disulfide between the β and γ chain. **(F)** Structure predictions for all LmE8 isoforms are high confidence and coiled-coils align well: LmE8-111 (pTM: 0.66, ipTM: 0.86), LmE8-511 (pTM: 0.88, ipTM: 0.87), LmE8-411 (pTM: 0.85, ipTM: 0.85), LmE8-211 (pTM: 0.75, ipTM: 0.84), LmE8-221 (pTM: 0.80, ipTM: 0.84), LmE8-523 (pTM: 0.88, ipTM: 0.87).

In a pilot study, LmE8-332 without the engineered disulfide and without the unstructured loop expressed with higher yield and more efficient heterotrimer formation than the original construct, with most of the protein forming the correct heterotrimer **(Figure D-E)**. Encouraged by these results, we re-designed all laminin chains and generated structure predictions in AlphaFold3 Multimer with the new design strategy. Inspection of the structure predictions confirmed that the coiled-coils aligned well in all isoforms and that the unstructured loops were removed **(Figure 4F)**.

### Optimized LmE8 construct design enables purification of all LmE8 isoforms

To assess if the redesigned LmE8 constructs were modular and capable of interchange to allow purification of all LmE8 isoforms, we tested the expression of Lm111, Lm211, Lm221, Lm332, Lm411, and Lm511 (the γ3 chain was excluded from this analysis because its C-terminal tail is truncated and does not bind integrins (22)). Each LmE8 isoform was readily expressed, and each yielded fractions of the correct heterotrimer size after affinity purification followed by size exclusion chromatograhy **(Figure 5A,D).** In each case, all three chains of the heterotrimer were present and formed the correct β-γ disulfide bond **(Figure 5B,E)**. The Lm211 and Lm221 peaks were slightly left shifted **(Figure 5D)** but had all heterotrimer components and formed the correct β-γ disulfide bond **(Figure 5E).**

**Figure 5.**
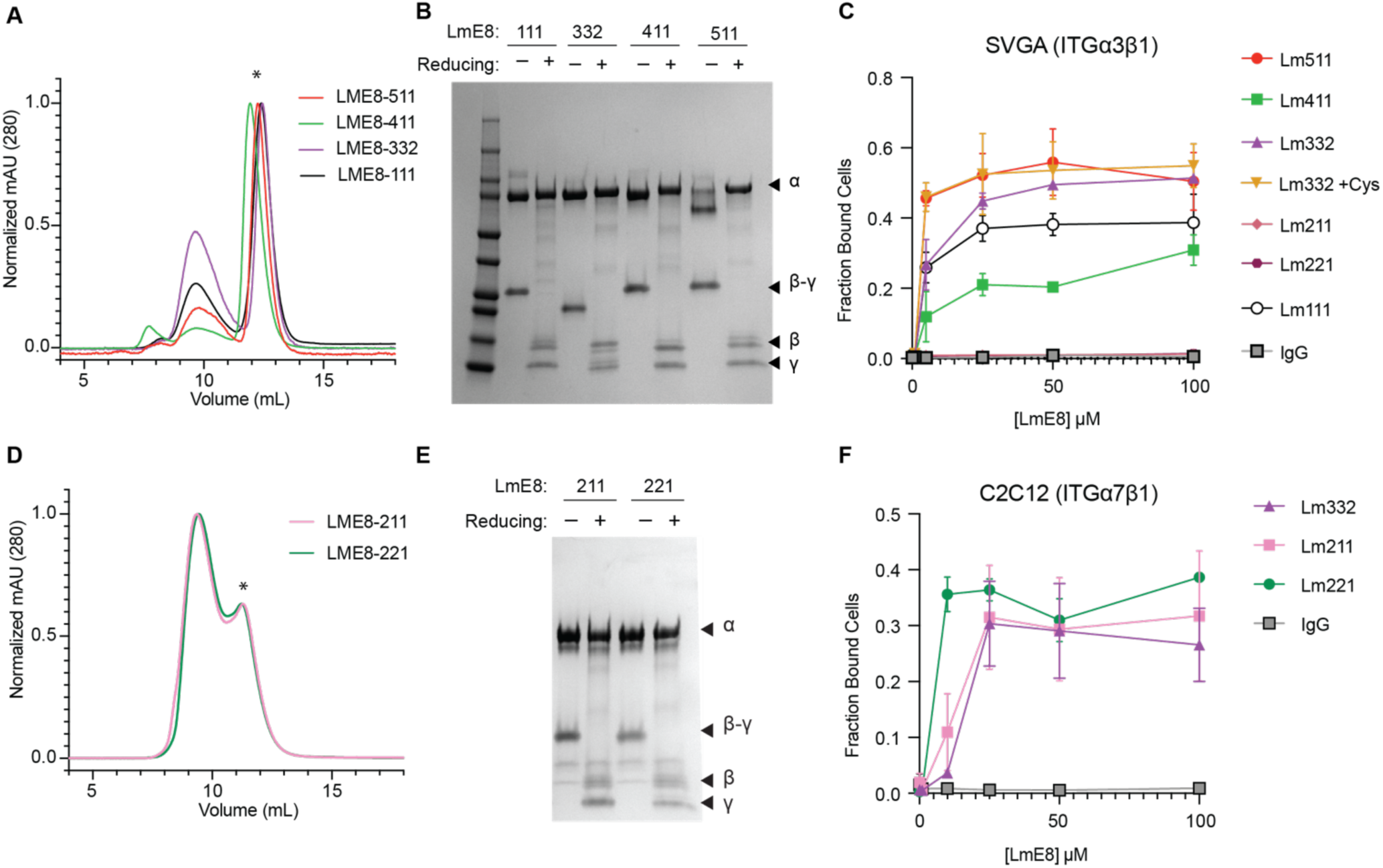
Expression and adhesion of the full panel of designed LmE8 proteins. (A,D) Size exclusion chromatograms (SEC) showing purifications of all LmE8 proteins on a Superdex 200i column. Traces were normalized to set the maximum and minimum 280 nm absorbance at 1 and 0, respectively. The heterotrimer peak is indicated with an asterisk. **(B,E)** Coomassie stained SDS-PAGE gels of SEC purified LmE8 proteins under non-reducing and reducing conditions, showing reduction of the native β-γ disulfide bond in the reducing sample. **(C,F)** SVG-A cells, which express ITGα3β1 and ITGα6β1 but not ITGα7β1, adhere to plates coated with all LmE8 isoforms except Lm211 and Lm221. C2C12 cells, which express ITGα7β1, adhere to Lm211 and Lm221 coated plates.

To confirm that these optimized LmE8 isoforms were also functional, we used our cell- based adhesion assay. SVG-A cells adhered to all LmE8 isoforms except Lm211 and Lm221 **(Figure 5C)**. This result is highly consistent with SVG-A cells having no ITGα7β1 expression (23), which is the only integrin with high affinity for Lm211 and Lm221 (13). Therefore, we also tested adhesion on LmE8 of C2C12 mouse myoblast cells which have high ITGα7β1 expression (24). C2C12 cells adhere to Lm211, Lm221, and Lm332 **(Figure 5F)**, and we thus conclude all the LmE8 fragments are functional.

## Discussion

In this study, we developed a modular, interchangeable approach for expression and purification of all integrin-binding laminins. Optimization guided by AlphaFold 3 predictions increased expression, improved heterotrimer formation, and enabled purification of all laminin isoforms. To make this approach broadly accessible to researchers, laminin chain sequences are provided in **(Table 1)** and plasmids will be deposited with Addgene.

Although LmE8-111, LmE8-221, LmE8-332, LmE8-411, and LmE8-511 fragments, are commercially available from iMatrix, our approach enables purification of all 17 biochemically confirmed laminin isoforms that are expressed throughout development and in various tissues (1,25,26). For example, Lm-511 is expressed during embryonic development and in all adult tissue basement membranes (27). Laminin α4-containing isoforms are present in the vascular endothelium basement membrane (28). Lm-332 is the major component of the skin basement membrane, and mutating any chain leads to junctional epidermolysis bullosa (29). Lm-211 is expressed in muscle fibers and α2 mutation leads to congenital muscular dystrophies (30). LmE8 fragments, therefore, could be useful reagents for studying laminopathy disease states.

This panel will be useful for studying laminin biology and adhesion receptor recognition. Moreover, the truncated LmE8 forms of the laminins should be more amenable to structural biology approaches. To date, the only laminin-integrin structure that has been reported is of LmE8-511 bound to ITGα6β1 (14). Our pipeline should enable structure determination of other LME8-integrin complexes, which would then reveal how laminin-integrin pairing specificity is encoded in the different laminin isoforms.

LmE8 ligands have also been used to direct stem cell growth and differentiation (31,32). Full length Lm-111 is the primary component in mouse derived Matrigel used to induce 3D organoid cultures (4). However, batch effects, contamination, and the need for xeno-free methods for stem-cell research places limits on the use of Matrigel in translational studies (33). Developing methods to embed LmE8 fragments in hydrogels has the potential to resolve these concerns (32). More generally, our panel of LmE8 ligands will provide useful reagents for a wide range of studies encompassing integrin biology, tissue growth in 3D cultures, and methods for organoid development and propagation.

## Materials and Methods

### Laminin E8 fragment design and cloning

Laminin chains were aligned using ClustalOmega, and initial construct length was determined by alignment to Lm511 crystal structure (14). Gene blocks for laminin α, β, and γ chains were codon optimized and all BsaI and BsmBI restriction sites were removed to allow for Golden Gate cloning. Laminin chain and signal sequence + affinity tag were cloned into part plasmids using BsmBI (NEB), enabling easy tag swaps. Final plasmids with an HA signal sequence, affinity tag, and laminin chain were cloned into the pTarget vector using BsaI (NEB). Cysteines were mutated to remove engineered disulfides using QuikChange mutagenesis kits (Aligent) with primers designed using the Aligent primer design program. Laminin chain truncations were PCR amplified using KOD1 polymerase (Sigma Aldrich) and cloned into vectors using BsaI (NEB).

### Laminin expression and purification

Constructs were transfected in Expi293T cells (Thermo Fisher) at a 1:1:1 (w:w:w) ratio with FectoPro (VWR) at 1 µg DNA:0.8 µl FectoPro per ml culture volume. Cells were enhanced 24 hrs after transfection with 3mM valproic acid and 0.4% D-(+)-glucose (Sigma-Aldrich). After viability dropped below 50%, approximately 5-6 days, cultures were spun down for 15 minutes at 4,000xg and supernatant was harvested. 1 mM CaCl2 was added to supernatants, which were then passed over 6 ml of M1-Flag resin (made in-house) to affinity purify the complex through the tag on the γ subunit. Resin was washed with 2x50 ml HBS (20 mM HEPES pH 7.5, 150 mM NaCl) + 1 mM CaCl2. Samples were eluted from the resin with HBS + 0.1 mg/ml Flag peptide + 5 mM EDTA. Elutions were concentrated in a 50 kDa concentrator (Millipore Sigma) and then loaded onto a Superose 6 increase or Superose 200 (Cytiva) increase size exclusion column to purify only LmE8 that contained all chains. Fractions were confirmed to have all laminin chains by reducing and non-reducing SDS-PAGE, and appropriate fractions were concentrated in a 50 kDa concentrator. Samples were flash frozen and stored at -80°C.

### Cell culture

SVG-A human fetal astrocyte cells were from Walter J. Atwood at Brown University. C2C12 mouse myoblast cells were purchased from ATCC. Both cell lines were cultured at 37 °C and 5% CO2 in DMEM (Corning) with 10% fetal bovine serum (FBS, GeminiBio) and 1% Penicillin- Streptomycin (PS, Thermo Fisher).

### Genomic engineering for ITGα3 and ITGα6 knockouts

ITGα3 and ITGα6 knockouts (KOs) were generated using the methodology described in Chapter 2. Guide RNAs (gRNAs) targeting ITGα3 and ITGα6 were designed using CRISPOR for high predicted efficiency and minimal off target effects. gRNAs were cloned into the PX458 Cas9/GFP plasmid (Addgene). SVG-A cells were transiently transfected using Lipofectamine 2000 (Thermo Fisher). 12 hours after transfection, cells were washed and fresh media was added. 2 days after transfection, cells were sorted using a Sony SH800 flow cytometer and cell sorter. GFP positive cells were single cell sorted into 96 well plates with 40% conditioned DMEM, 40% fresh DMEM, and 20% FBS. Single cell clones were screened by western blot, flow cytometry, and/or genomic DNA PCR fragment sequencing. Using this approach, we generated ITGα3, ITGα6, and ITGα3/ITGα6 KO clones to assess cell adhesion to LmE8 fragments.

### Cell adhesion assay

LmE8 fragments were immobilized on 96 well high binding black/clear bottom plates (Corning) by adding 100 µl LmE8 at varying concentrations and incubating at 4 °C overnight. The next day, plates were washed twice with PBS and blocked with PBS containing 30 mg/ml BSA (Roche) at 37 °C for 1 hr. Plates were then washed twice with PBS. SVG-A KO cells were trypsinized for 5 min with Trypsin/0.53 mM EDTA (Corning) and diluted to 2.0 x 10^5^ cells/mL in DMEM without phenol red containing 1 µM BCECF AM dye (Thermo Fisher). 100 µl of diluted cells were added to each well and allowed to adhere for 1 hr at 37 °C in an incubator containing 5% CO2. Cells were washed 4 times with a BioTEK plate washer and resuspended in PBS. BCECF AM fluorescence was measured on a Spectra Max analyzer using excitation at 440 nm for emission at 535 nm. Background signal was subtracted and the fraction of cells bound before and after washing was calculated.

To assess FAK activation, 150 nM of the LmE8 fragment was immobilized on 6 well untreated tissue culture plates (Corning). Plates were then prepared as previously described. 200,000 SVG-A cells were added per well and spun down for 1 min at 100 g. At each timepoint, media was aspirated and 200 µl of SDS loading dye containing 1x PhosStop inhibitor (Sigma- Aldrich) was added. For Western blots, 10 µl of each sample was run on a gel and transferred to a nitrocellulose membrane (VWR). Blots were blocked with 5% BSA in TBST (20 mM Tris pH 7.5, 150 mM NaCl, 0.1% Tween 20). Blots were incubated with GAPDH and P-FAK primary antibodies (Cell Signaling Technologies) in TBST containing 1% BSA overnight at 4 °C per manufacturer instructions. HRP conjugated secondary antibodies (Cell Signaling Technologies) were applied for 1 hr at room temperature. Blots were washed with TBST and imaged.

### AlphaFold 3 multimer structure predictions

AlphaFold 3 multimer structure predictions were generated with default settings (17). Confidence statistics (ipTM, pTM, and PAE plots) were generated from the server. Predicted Alignment Error (Å) (PAE) is a confidence metric between two residues. Lower values indicate lower predicted error and higher prediction confidence. Predicted template modeling (pTM) and interface predicted template modeling (ipTM) are additional confidence metrics that predict the accuracy of the whole structure. AlphaFold 3 defines high-quality predictions above 0.8. Structure predictions were visualized in PyMol.

## Supplementary material description

**Table 1.** Laminin chain sequences

| Laminin chain | Protein sequence |
|---|---|
| α1 | TTLRETHQLLQDSTMATLLAGRKVKDVEIQANLLFDRLKPLKMLEENLSRNLSEIKLLISQARKQAASIKVAVSADRDCIRAYQPQISSTNYNTLTLNVKTQEPDNLLFYLGSSTASDFLAVEMRRGRVAFLWDLGSGSTRLEFPDFPIDDNRWHSIHVARFGNIGSLSVKEMSSNQKSPTKTSKSPGTANVLDVNNSTLMFVGGLGGQIKKSPAVKVTHFKGCLGEAFLNGKSIGLWNYIEREGKCRGCFGSSQNEDPSFHFDGSGYSVVEKSLPATVTQIIMLFNTFSPNGLLLYLGSYGTKDFLSIELFRGRVKVMTDLGSGPITLLTDRRYNNGTWYKIAFQRNRKQGVLAVIDAYNTSNKETKQGETPGASSDLNRLDKDPIYVGGLPRSRVVRRGVTTKSFVGCIKNLEISRSTFDLLRNSYGVRKGCLLEPIRSVSFLKGGYIELPPKSLSPESEWLVTFATTNSSGIILAALGGDVEKRGDREEAHVPFFSVMLIGGNIEVHVNPGDGTGLRKALLHAPTGTCSDGQAHSISLVRNRRIITVQLDENNPVEMKLGTLVESRTINVSNLYVGGIPEGEGTSLLTMRRSFHGCIKNLIFNLELLDFNSAVGHEQVDLDTCWLSERP |
| α2 | DSVAKTNAVVKDPSKNKIIADADATVKNLEQEADRLIDKLKPIKELEDNLKKNISEIKELINQARKQANSIKVSVSSGGDCIRTYKPEIKKGSYNNIVVNVKTAVADNLLFYLGSAKFIDFLAIEMRKGKVSFLWDVGSGVGRVEYPDLTIDDSYWYRIVASRTGRNGTISVRALDGPKASIVPSTHHSTSPPGYTILDVDANAMLFVGGLTGKLKKADAVRVITFTGCMGETYFDNKPIGLWNFREKEGDCKGCTVSPQVEDSEGTIQFDGEGYALVSRPIRWYPNISTVMFKFRTFSSSALLMYLATRDLRDFMSVELTDGHIKVSYDLGSGMASVVSNQNHNDGKWKSFTLSRIQKQANISIVDIDTNQEENIATSSSGNNFGLDLKADDKIYFGGLPTLRNLSMKARPEVNLKKYSGCLKDIEISRTPYNILSSPDYVGVTKGCSLENVYTVSFPKPGFVELSPVPIDVGTEINLSFSTKNESGIILLGSGGTPAPPRRKRRQTGQAYYAILLNRGRLEVHLSTGARTMRKIVIRPEPNLFHDGREHSVHVERTRGIFTVQVDENRRYMQNLTVEQPIEVKKLFVGGAPPEFQPSPLRNIPPFEGCIWNLVINSVPMDFARPVSFKNADIGRCAHQKLR |
| α3 | NEDFKKALTDADNSVNKLTNKLPDLWRKIESINQQLLPLGNISDNMDRIRELIQQARDAASKVAVPMRFNGKSGVEVRLPNDLEDLKGYTSLSLFLQRPNSRENGGTENMFVMYLGNKDASRDYIGMAVVDGQLTCVYNLGDREAELQVDQILTKSETKEAVMDRVKFQRIYQFARLNYTKGATSSKPETPGVYDMDGRNSNTLLNLDPENVVFYVGGYPPDFKLPSRLSFPPYKGCIELDDLNENVLSLYNFKKTFNLNTTEVEPCRRRKEESDKNYFEGTGYARVPTQPHAPIPTFGQTIQTTVDRGLLFFAENGDRFISLNIEDGKLMVRYKLNSELPKERGVGDAINNGRDHSIQIKIGKLQKRMWINVDVQNTIIDGEVFDFSTYYLGGIPIAIRERFNISTPAFRGCMKNLKKTSGVVRLNDTVGVTKKCSEDWKLVRSASFSRGGQLSFTDLGLPPTDHLQASFGFQTFQPSGILLDHQTWTRNLQVTLEDGYIELSTSDSGGPIFKSPQTYMDGLLHYVSVISDNSGLRLLIDDQLLRNSKRLKHISSSRQSLRLGGSNFEGCISNVFVQRLSLSPEVLDLTSNSLKRDVSLGGCSLNKPPFLMLLKGS |
| α4 | SSAYNTAVNSARDAVRNLTEVVPQLLDQLRTVEQKRPASNVSASIQRIRELIAQTRSVASKIQVSMMFDGQSAVEVHSRTSMDDLKAFTSLSLYMKPPVKRPELTETADQFILYLGSKNAKKEYMGLAIKNDNLVYVYNLGTKDVEIPLDSKPVSSWPAYFSIVKIERVGKHGKVFLTVPSLSSTAEEKFIKKGEFSGDDSLLDLDPEDTVFYVGGVPSNFKLPTSLNLPGFVGCLELATLNNDVISLYNFKHIYNMDPSTSVPCARDKLAFTQSRAASYFFDGSGYAVVRDITRRGKFGQVTRFDIEVRTPADNGLILLMVNGSMFFRLEMRNGYLHVFYDFGFSGGPVHLEDTLKKAQINDAKYHEISIIYHNDKKMILVVDRRHVKSMDNEKMKIPFTDIYIGGAPPEILQSRALRAHLPLDINFRGCMKGFQFQKKDFNLLEQTETLGVGYGCPEDSLISRRAYFNGQSFIASIQKISFFDGFEGGFNFRTLQPNGLLFYYASGSDVFSISLDNGTVIMDVKGIKVQSVDKQYNDGLSHFVISSVSPTRYELIVDKSRVGSKNPTKGKIEQTQASEKKFYFGGSPISAQYANFTGCISNAYFTRVDRDVEVEDFQRYTEKVHTSLYECPIESS |
| α5 | GQDLGQAVLDAGHSVSTLEKTLPQLLAKLSILENRGVHNASLALSASIGRVRELIAQARGAASKVKVPMKFNGRSGVQLRTPRDLADLAAYTALKFYLQGPEPEPGQGTEDRFVMYMGSRQATGDYMGVSLRDKKVHWVYQLGEAGPAVLSIDEDIGEQFAAVSLDRTLQFGHMSVTVERQMIQETKGDTVAPGAEGLLNLRPDDFVFYVGGYPSTFTPPPLLRFPGYRGCIEMDTLNEEVVSLYNFERTFQLDTAVDRPCARSKSTGDPWLTDGSYLDGTGFARISFDSQISTTKRFEQELRLVSYSGVLFFLKQQSQFLCLAVQEGSLVLLYDFGAGLKKAVPLQPPPPLTSASKAIQVFLLGGSRKRVLVRVERATVYSVEQDNDLELADAYYLGGVPPDQLPPSLRRLFPTGGSVRGCVKGIKALGKYVDLKRLNTTGVSAGCTADLLVGRAMTFHGHGFLRLALSNVAPLTGNVYSGFGFHSAQDSALLYYRASPDGLCQVSLQQGRVSLQLLRTEVKTQAGFADGAPHYVAFYSNATGVWLYVDDQLQQMKPHRGPPPELQPQPEGPPRLLLGGLPESGTIYNFSGCISNVFVQRLLGPQRVFDLQQNLGSVNVSTGCAPALQAQ |
| β1 | DARRKAEMLQNEAKTLLAQANSKLQLLKDLERKYEDNQRYLEDKAQELARLEGEVRSLLKDISQKVAVYSTCL |
| β2 | AAQARAEQLRDEARDLLQAAQDKLQRLQELEGTYEENERALESKAAQLDGLEARMRSVLQAINLQVQIYNTCQ |
| β3 | EQGARIQSVKTEAEELFGETMEMMDRMKDMELELLRGSQAIMLRSADLTGLEKRVEQIRDHINGRVLYYATCK |
| γ1 | GDTVDLNKLNEIEGTLNKAKDEMKVSDLDRKVSDLENEAKKQEAAIMDYNRDIEEIMKDIRNLEDIRKTLPSGCFNTPSIEKP |
| γ2 | GLSVDEEGLVLLEQKLSRAKTQINSQLRPMMSELEERARQQRGHLHLLETSIDGILADVKNLENIRDNLPPGCYNTQALEQQ |
| γ3 | GHQAPAQALNETQWALERLRLQLGSPGSLQRKLSLLEQESQQQELQIQGFESDLAEIRADKQNLEAILHSLPENCASWQ |

## Acknowledgements

Financial support for this work was provided by NIH grant R01 AI172846 (to S.C.B.), a gift from Edward B. Goodnow (to S.C.B.), and F31 AR083740 to LL. We thank Yuxin Hao and Tim Springer’s laboratory for advice on cell adhesion assay protocols. We also thank members of the Blacklow and Kruse labs for helpful discussions.

## Data Availability Statement

The data that support the findings of this study are available from the corresponding authors upon reasonable request.

## AI Disclosure Statement

AI was not used in idea generation, data interpretation, or preparation of this manuscript.

## Notes

### Competing Interest Statement

The authors have declared no competing interest.

## References

1. Colognato H, Yurchenco PD. Form and function: the laminin family of heterotrimers. Dev Dyn. 2000 Jun;218(2):213–34. doi:10.1002/(SICI)1097-0177(200006)218:2<213::AID-DVDY1>3.0.CO;2-R PubMed PMID: 10842354.

2. Martin GR, Timpl R. Laminin and other basement membrane components. Annu Rev Cell Biol. 1987;3:57–85. doi:10.1146/annurev.cb.03.110187.000421 PubMed PMID: 3318883.

3. Yao Y. Laminin: loss-of-function studies. Cell Mol Life Sci. 2017 Mar;74(6):1095–115. doi:10.1007/s00018-016-2381-0 PubMed PMID: 27696112; PubMed Central PMCID: PMC11107706.

4. Clevers H. Modeling Development and Disease with Organoids. Cell. 2016 Jun 16;165(7):1586–97. doi:10.1016/j.cell.2016.05.082 PubMed PMID: 27315476.

5. Aumailley M, Bruckner-Tuderman L, Carter WG, Deutzmann R, Edgar D, Ekblom P, et al. A simplified laminin nomenclature. Matrix Biol. 2005 Aug;24(5):326–32. doi:10.1016/j.matbio.2005.05.006 PubMed PMID: 15979864.

6. Engel J, Odermatt E, Engel A, Madri JA, Furthmayr H, Rohde H, et al. Shapes, domain organizations and flexibility of laminin and fibronectin, two multifunctional proteins of the extracellular matrix. J Mol Biol. 1981 Jul 25;150(1):97–120. doi:10.1016/0022-2836(81)90326-0 PubMed PMID: 6795355.

7. Hohenester E. Structural biology of laminins. Essays Biochem. 2019 Sep 13;63(3):285–95. doi:10.1042/EBC20180075 PubMed PMID: 31092689; PubMed Central PMCID: PMC6744579.

8. Hussain SA, Carafoli F, Hohenester E. Determinants of laminin polymerization revealed by the structure of the α5 chain amino-terminal region. EMBO Rep. 2011 Mar;12(3):276–82. doi:10.1038/embor.2011.3 PubMed PMID: 21311558; PubMed Central PMCID: PMC3059903.

9. Kulczyk AW, McKee KK, Zhang X, Bizukojc I, Yu YQ, Yurchenco PD. Cryo-EM reveals the molecular basis oflaminin polymerization and LN-lamininopathies. Nat Commun. 2023 Jan 19;14(1):317. doi:10.1038/s41467-023-36077-z PubMed PMID: 36658135; PubMed Central PMCID: PMC9852560.

10. Denzer AJ, Brandenberger R, Gesemann M, Chiquet M, Ruegg MA. Agrin binds to the nerve-muscle basal lamina via laminin. J Cell Biol. 1997 May 5;137(3):671–83. doi:10.1083/jcb.137.3.671 PubMed PMID: 9151673; PubMed Central PMCID: PMC2139873.

11. Takagi J, Yang Y, Liu JH, Wang JH, Springer TA. Complex between nidogen and laminin fragments reveals a paradigmatic beta-propeller interface. Nature. 2003 Aug 21;424(6951):969–74. doi:10.1038/nature01873 PubMed PMID: 12931195.

12. Goodman SL, Risse G, von der Mark K. The E8 subfragment of laminin promotes locomotion of myoblasts over extracellular matrix. J Cell Biol. 1989 Aug;109(2):799–809. doi:10.1083/jcb.109.2.799 PubMed PMID: 2503526; PubMed Central PMCID: PMC2115702.

13. Nishiuchi R, Takagi J, Hayashi M, Ido H, Yagi Y, Sanzen N, et al. Ligand-binding specificities of laminin-binding integrins: a comprehensive survey of laminin-integrin interactions using recombinant alpha3beta1, alpha6beta1, alpha7beta1 and alpha6beta4 integrins. Matrix Biol. 2006 Apr;25(3):189–97. doi:10.1016/j.matbio.2005.12.001 PubMed PMID: 16413178.

14. Takizawa M, Arimori T, Taniguchi Y, Kitago Y, Yamashita E, Takagi J, et al. Mechanistic basis for the recognition of laminin-511 by α6β1 integrin. Sci Adv. 2017 Sep;3(9):e1701497. doi:10.1126/sciadv.1701497 PubMed PMID: 28879238; PubMed Central PMCID: PMC5580876.

15. Pulido D, Hussain SA, Hohenester E. Crystal Structure of the Heterotrimeric Integrin- Binding Region of Laminin-111. Structure. 2017 Mar 7;25(3):530–5. doi:10.1016/j.str.2017.01.002 PubMed PMID: 28132784; PubMed Central PMCID: PMC5343747.

16. Ido H, Nakamura A, Kobayashi R, Ito S, Li S, Futaki S, et al. The requirement of the glutamic acid residue at the third position from the carboxyl termini of the laminin gamma chains in integrin binding by laminins. J Biol Chem. 2007 Apr 13;282(15):11144–54. doi:10.1074/jbc.M609402200 PubMed PMID: 17307733.

17. Abramson J, Adler J, Dunger J, Evans R, Green T, Pritzel A, et al. Accurate structure prediction of biomolecular interactions with AlphaFold 3. Nature. 2024 Jun;630(8016):493–500. doi:10.1038/s41586-024-07487-w PubMed PMID: 38718835; PubMed Central PMCID: PMC11168924.

18. Saito-Diaz K, Saini T, Patel AJ, James C, Thomas KS, Knight TN, et al. Laminin β4 is required for the development of human peripheral sensory neurons. bioRxiv: The Preprint Server for Biology; 2024. doi:10.1101/2024.11.22.624899 PubMed PMID: 39605577; PubMed Central PMCID: PMC11601542.

19. Aumailley M. The laminin family. Cell Adh Migr. 2013;7(1):48–55. doi:10.4161/cam.22826 PubMed PMID: 23263632; PubMed Central PMCID: PMC3544786.

20. Lietha D, Cai X, Ceccarelli DFJ, Li Y, Schaller MD, Eck MJ. Structural basis for the autoinhibition of focal adhesion kinase. Cell. 2007 Jun 15;129(6):1177–87. doi:10.1016/j.cell.2007.05.041 PubMed PMID: 17574028; PubMed Central PMCID: PMC2077847.

21. Yazlovitskaya EM, Viquez OM, Tu T, De Arcangelis A, Georges-Labouesse E, Sonnenberg A, et al. The laminin binding α3 and α6 integrins cooperate to promote epithelial cell adhesion and growth. Matrix Biol. 2019 Apr;77:101–16. doi:10.1016/j.matbio.2018.08.010 PubMed PMID: 30193894; PubMed Central PMCID: PMC6399080.

22. Ido H, Ito S, Taniguchi Y, Hayashi M, Sato-Nishiuchi R, Sanzen N, et al. Laminin isoforms containing the gamma3 chain are unable to bind to integrins due to the absence of the glutamic acid residue conserved in the C-terminal regions of the gamma1 and gamma2 chains. J Biol Chem. 2008 Oct 17;283(42):28149–57. doi:10.1074/jbc.M803553200 PubMed PMID: 18697739; PubMed Central PMCID: PMC2661386.

23. Tveriakhina L, Scanavachi G, Egan ED, Da Cunha Correia RB, Martin AP, Rogers JM, et al. Temporal dynamics and stoichiometry in human Notch signaling from Notch synaptic complex formation to nuclear entry of the Notch intracellular domain. Dev Cell. 2024 Jun 3;59(11):1425–1438.e8. doi:10.1016/j.devcel.2024.03.021 PubMed PMID: 38574735; PubMed Central PMCID: PMC11153029.

24. Zolkiewska A, Thompson WC, Moss J. Interaction of integrin alpha 7 beta 1 in C2C12 myotubes and in solution with laminin. Exp Cell Res. 1998 Apr 10;240(1):86–94. doi:10.1006/excr.1998.4002 PubMed PMID: 9570924.

25. Domogatskaya A, Rodin S, Tryggvason K. Functional diversity of laminins. Annu Rev Cell Dev Biol. 2012;28:523–53. doi:10.1146/annurev-cellbio-101011-155750 PubMed PMID: 23057746.

26. Yap L, Tay HG, Nguyen MTX, Tjin MS, Tryggvason K. Laminins in Cellular Differentiation. Trends in Cell Biology. 2019 Dec;29(12):987–1000. doi:10.1016/j.tcb.2019.10.001

27. Miner JH, Patton BL, Lentz SI, Gilbert DJ, Snider WD, Jenkins NA, et al. The laminin alpha chains: expression, developmental transitions, and chromosomal locations of alpha1-5, identification of heterotrimeric laminins 8-11, and cloning of a novel alpha3 isoform. J Cell Biol. 1997 May 5;137(3):685–701. doi:10.1083/jcb.137.3.685 PubMed PMID: 9151674; PubMed Central PMCID: PMC2139892.

28. Wang J, Hoshijima M, Lam J, Zhou Z, Jokiel A, Dalton ND, et al. Cardiomyopathy associated with microcirculation dysfunction in laminin alpha4 chain-deficient mice. J Biol Chem. 2006 Jan 6;281(1):213–20. doi:10.1074/jbc.M505061200 PubMed PMID: 16204254.

29. Kiritsi D, Has C, Bruckner-Tuderman L. Laminin 332 in junctional epidermolysis bullosa. Cell Adh Migr. 2013;7(1):135–41. doi:10.4161/cam.22418 PubMed PMID: 23076207; PubMed Central PMCID: PMC3544777.

30. Sinha R, Singh S, MONA TIWARI, ZULFIKAR LUHAR. The evolving spectrum of LAMA2 related congenital muscular dystrophy (MDC1) [Internet]. 2024. doi:10.13140/RG.2.2.28044.53127

31. Ohta R, Niwa A, Taniguchi Y, Suzuki NM, Toga J, Yagi E, et al. Laminin-guided highly efficient endothelial commitment from human pluripotent stem cells. Sci Rep. 2016 Nov 2;6(1):35680. doi:10.1038/srep35680

32. Miyazaki T, Futaki S, Suemori H, Taniguchi Y, Yamada M, Kawasaki M, et al. Laminin E8 fragments support efficient adhesion and expansion of dissociated human pluripotent stem cells. Nat Commun. 2012;3:1236. doi:10.1038/ncomms2231 PubMed PMID: 23212365; PubMed Central PMCID: PMC3535336.

33. Wolff L, Hendrix S. Rethinking Matrigel: The Complex Journey to Matrix Alternatives in Organoid Culture. Adv Sci (Weinh). 2025 Dec;12(47):e08734. doi:10.1002/advs.202508734 PubMed PMID: 41204762; PubMed Central PMCID: PMC12713094.

